## Supplementary file for "Cryo-EM Structures of Higher Order Gephyrin Oligomers Reveal Principles of Inhibitory Postsynaptic Scaffold Organization"

Author to whom correspondence should be addressed:

### Supplementary Material

#### Cloning and mutagenesis

Gephyrin mScarlet variants R379E, D422N, and $^{314}$EEHE$^{317}$ (R314E, R315E, R317E) were generated by site-directed mutagenesis PCR from the construct GephFL-mScarlet, encoding Gephyrin P1 splicing variant fused to mScarlet [1]. The gene encoding GephE$_{309}$ encompassing residues 309-726 was amplified by PCR and cloned using ligation into the pET-28b plasmid using the NheI and HindIII restriction enzymes sites. A list of all constructs used in this work is available in Table S1.

#### CryoEM data processing for the GephE-27F3 complex and structure solution

Template picked particles were extracted to a box size of 320 x 320 and cropped to 240 x 240 pixels (pixel size of 2.92 Å/pix) and screened by 2D classification in CryoSPARC [2]. After ab-initio reconstruction and sequential heterogenous refinement, 1,670,313 particles from the best classes were combined with 3,613,232 particles from Topaz picking and 2D classification and duplicates were removed. The joined particles were classified again using 2D classification and 2,166,682 particles from selected classes were used for an ab-initio reconstruction with 4 classes. The particles were subjected to heterogeneous refinement and the best class with 2,166,682 particles was used for a subsequent 3D classification. The best class with 1,233,122 particles was subjected to non-uniformity (NU) refinement to produce a volume with a resolution of 2.4 Å at a pixel size of 0.97 Å/pix. 3D classification with 2 classes led to a final class with 1,233,122 particles that was NU-refined to a resolution of 2.4 Å. Extraction of the particles to a box size of 288 x 288 pixels and further NU-refinement resulted in a map with 780,576 particles and a resolution of 2.31 Å. All steps are summarized in Fig. S2.

#### CryoEM data processing for the GephFL-27F3 complex

Template picked particles were extracted to a box size of 320 x 320 and cropped to 92 x 92 pixels (pixel size of 2.92 Å/pix) and screened by 2D classification in CryoSPARC [2]. The resulting 963,369 particles were used for ab-initio construction with 10 classes. After heterogenous refinement, particles from the best class were extracted to a box size of 320 x 320 pixels at a pixel size of 0.84 Å/pix and refined using NU refinement resulting in a resolution of 3.2 Å. Particles obtained from Topaz picking and subsequent 2D classification were joined with the refined particles down sampled to a box size of 92 x 92 pix (2.8 Å/pix) and duplicates were removed. 718,277 particles were NU-refined and 3D variability analysis was used to generate 5 clusters. The 5 clusters were subjected to heterogenous refinement and the best class with 186‘910 particles was selected. In this cluster, the presence of extra-densities in comparison with the rest of the clusters was evident. The particles from this class were extracted to a box size of 320 x 320 pixels at a pixel size of 0.84 Å/pix and were NU refine producing a with a resolution of 3.1 Å. After global CTF refinement and NU-refinement, the 3D variability job was used to create 3 clusters and the final map consisting of 62,978 particles had a resolution of 3.35 Å. All steps are summarized in Fig. S3.

#### CryoEM data processing for the GephFL dimer-of-dimers structure

For the dimer-of-dimers map, multiple picking strategies were employed including blob picking, template picking and Topaz picking [2, 3]. Particles were extracted to a box size of 256 x 256 pixels (at a pixel size of 1.55 Å/pix) and then subjected to several rounds of 2D classification for each picking strategy. Classes were selected based on well centered particles that represented a tetramer leaving out hexamers and dimers. Selected particles from each picking strategy were combined and duplicates were removed. 763,062 particles remained and were extracted to a box size of 240 x 240 pixels at a pixel size of 1.41 Å/pix. After another round of 2D classification 406,886 particles remained and were refined using non-uniform refine in Cryosparc producing a map with a nominal resolution of 3.3 Å. Using PyEM [4] the data were transferred to star format for use in RELION [5]. 3D Auto-Refine in RELION with BLUSH [6] led to a 3.3 Å map. Unaligned 3D classification produced 3 classes of which the best resolved class with 283,182 particles was chosen and further refined to a map of 3.5 Å. Local masks surrounding each dimer domain were created (SI Fig. S4) and each domain was locally refined resulting in maps of 3.07 Å for one dimer and 3.1 Å for the other dimer.

#### Table S1: Plasmids and constructs.

| Construct name | Encoded nrotein | Residues | Tags | Vector | Source | Experimental Use |
|---|---|---|---|---|---|---|
| GephE | Geph E-domain | 318-736 | Chitin-binding, intein fused. | pTWIN2 | Ref. [7] | DSC, CD spectroscopy |
| $GephE_{309}$ | Geph E-domain | 309-736 | N-terminal $His_6$ | pET-28b | This study | Cryo-EM, DSC, CD spectroscopy |
| GephFL (P1) | Full-length gephyrin | 1-736 (P1 splice variant) | N-terminal $His_6$ | pET-28a | Ref. [8] | Cryo-EM, SEC-MALS |
| DARPin 27F3 | DARPin 27F3 | - | N-terminal $His_6$ | pET-28a | Ref. [8] | Cryo-EM, binding assays |
| GephFL mScarlet | Full-length gephyrin | 1-736 (WT) | C-terminal mScarlet | pLVX | Ref. [1] | Cellular clustering assays (HEK293) |
| GephFL mScarlet R379E | Full-length gephyrin R379E mutant | 1-736 | C-terminal mScarlet | pLVX | This study | Cellular clustering assays |
| GephFL mScarlet D422N | Full-length gephyrin D422N mutant | 1-736 | C-terminal mScarlet | pLVX | This study | Cellular clustering assays |
| GephFL mscarlet $^{314}$EEHE$^{317}$ | Full-length gephyrin (RRHR→EEHE) | 1-736 | C-terminal mScarlet | pLVX | This study | Cellular clustering assays |
| GephFL (P2) | Full-length Gephyrin | 1-750 (P2 splice variant) | N-terminal $His_6$ | pET-28b | Ref. [9] | Control in comparative oligomerization analysis |

**Table S2: Data collection and refinement statistics.**

| | **GephE$_{309}$-27F3** | **GephFL-27F3** | **GephFL$_{dimer-of-dimers}$** |
|---|---|---|---|
| **Data collection and processing** | | | |
| Accelerating voltage (kV) | 300 | 300 | 300 |
| Magnification(x10$^3$) | 165 | 105 | 130 |
| Number of Movies | 26204 | 25183 | 27072 |
| Pixel size (Å) | 0.73 | 0.84 | 0.66 |
| Dose e$^-$/Å$^2$ | 44 | 45 | 42.6 |
| Box lengths (Å) | 70.26, 93.99, 135.96 | 68.88, 94.08, 137.76 | 71.81, 85.89, 229.50 |
| Box angles (°) | 90.00, 90.00, 90.00 | 90.00, 90.00, 90.00 | 90.00, 90.00, 90.00 |
| CryoSPARC/RELION resolution (Å) @FSC=0.143 | 2.3 | 3.4 | 3.1 |
| **Refinement** | | | |
| Model resolution (Å) @FSC=0.5 | 2.5 | 3.5 | 3.3 |
| CC mask | 0.82 | 0.81 | 0.71 |
| Chains | 3 | 3 | 4 |
| Protein residues | 967 | 1011 | 1672 |
| Atoms (H count) | 14784 (7422 H) | 15503 (7811 H) | 12749 (0 H) |
| RMSD in bonds lengths (Å) / angles (°) | 0.004 Å / 0.698° | 0.003 Å / 0.651° | 0.004 Å / 0.692° |
| Validation | | | |
| MolProbity score | 1.38 | 1.57 | 2.37 |
| Clash score | 6.22 | 7.24 | 11.85 |
| Rotamer outliers (%) | 0.86 | 0.71 | 6.9 |
| CaBLAM outliers (%) | 1.78 | 1.60 | 1.45 |
| Ramachandran statistics (%) | | | |
| Favored | 97.81 | 97.01 | 97.24 |
| Allowed | 2.19 | 2.99 | 2.76 |

| | | | |
|---|---|---|---|
| Outliers | 0.00 | 0.00 | 0.0 |

**Fig. S1: GephFL architecture and GephE subdomains.**

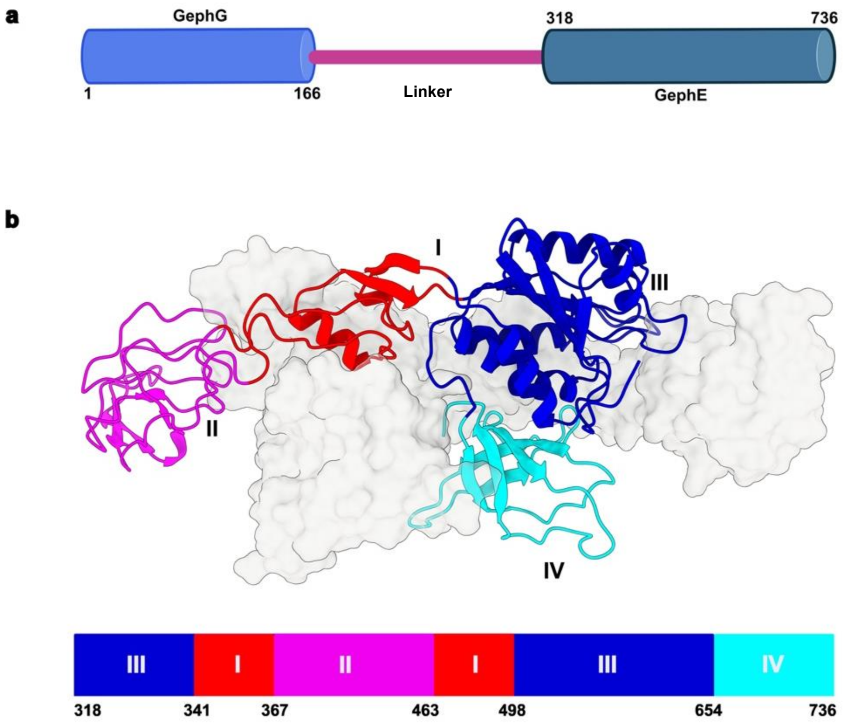


**a** Domain architecture of GephFL with N-terminal G-domain (GephG), central linker and C-terminal E-domain (GephE). **b** Structure of the GephE dimer with one subunit in ribbon representation, color coded to reflect its subdomain (I-IV) architecture and the other subunit in surface representation in gray. The color bar at the bottom defines the domain boundaries.

**Fig. S2: GephFL purification: Characterization of fraction A4 and SDS-PAGE analysis of different fractions.**

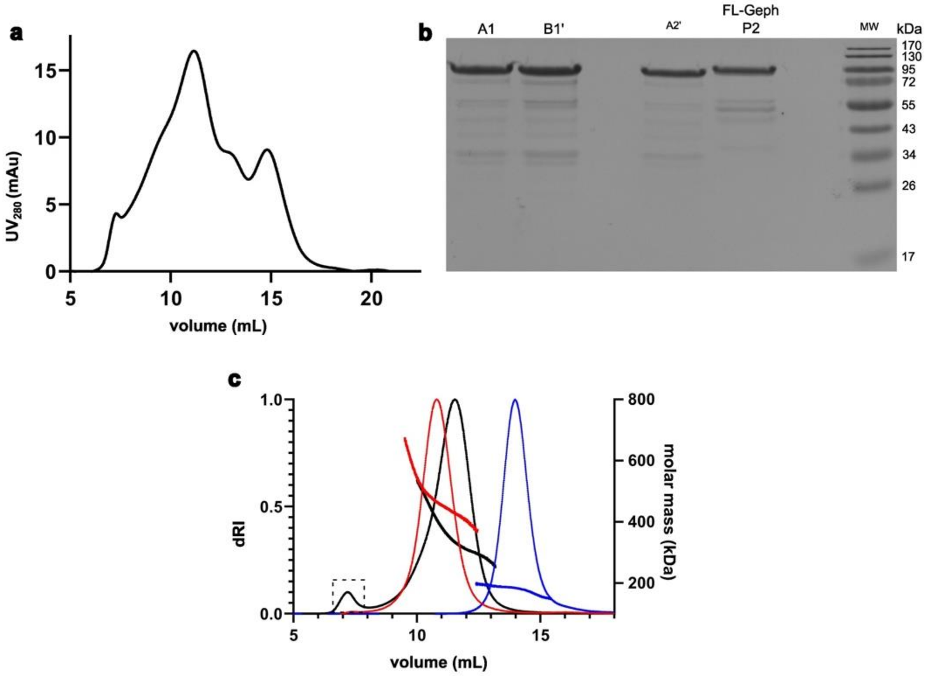


(a) Size exclusion chromatogram of GephFL fraction A4. Several peaks are present and hence this fraction was not further characterized. (b) SDS-PAGE of fractions A1, A2' and B1' of P1 GephFL (molecular weight of 82 kDa) and P2 GephFL P2 (molecular weight of 90 kDa). Bands at lower molecular weights represent degradation products which are commonly observed after gephyrin purification. The GephFL P2 was further analyzed in Fig. S8. (c) SEC MALS analyses of fractions A1 (red), A2' (black) and B1' (blue) across the entire chromatograms. A higher molecular weight species in A2' is highlighted by a dashed rectangle.

**Fig. S3: CryoEM data processing workflow for the GephE-27F3 complex.**

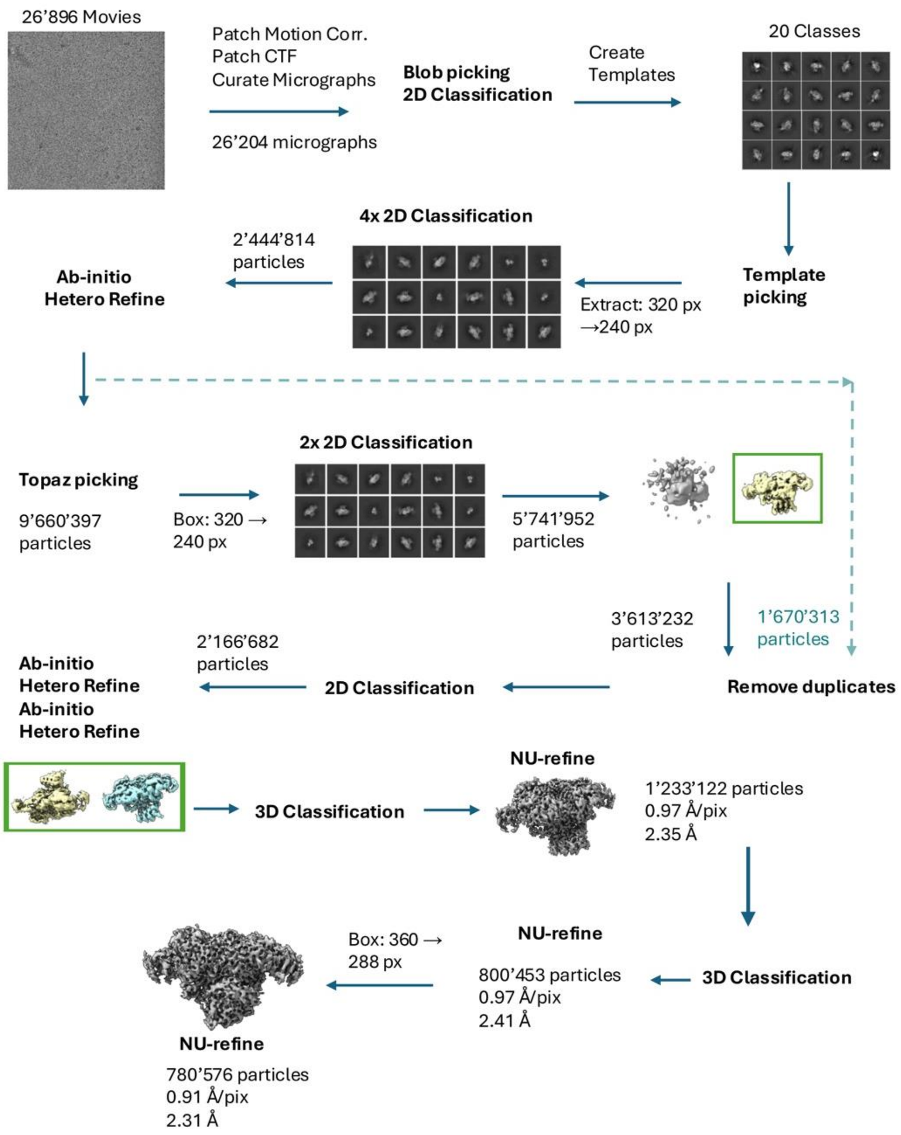

**Fig. S4: Superposition between GephFL-27F3 and the GephE-GlyR b receptor peptide.**

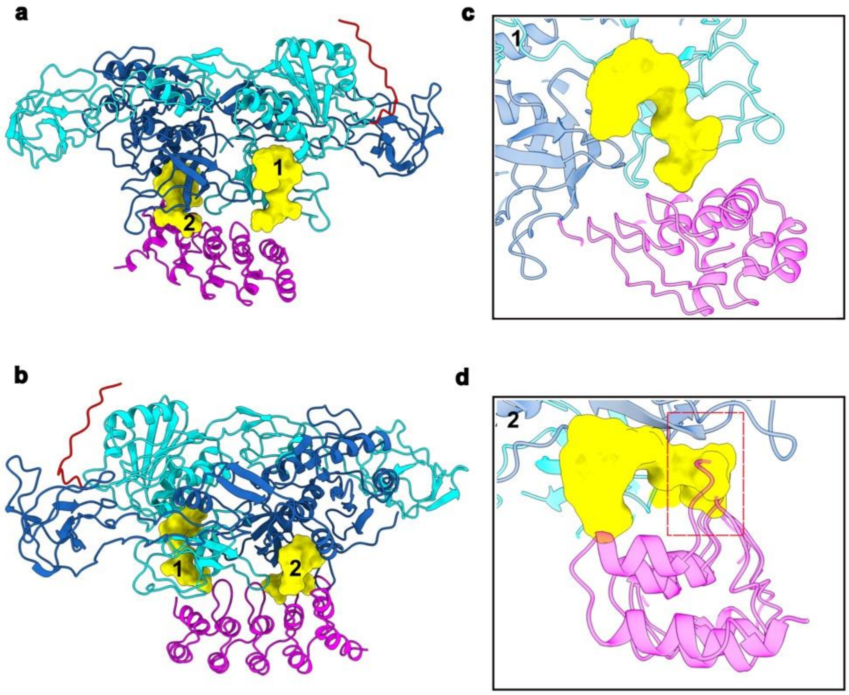


**a, b** Two orientations of the GephFL-27F3 complex in ribbon representation color coded as described in the main manuscript. The two GlyR b-subunit derived peptides (PDB entry 4PD1) are shown in surface representation in yellow. **c, d** Zoom into the binding sites. Binding site 1 can be occupied in the presence of 27F3 while binding of the peptide to site 2 is not compatible with 27F3 binding.

**Fig. S5: CryoEM data processing workflow for the GephFL-27F3 complex.**

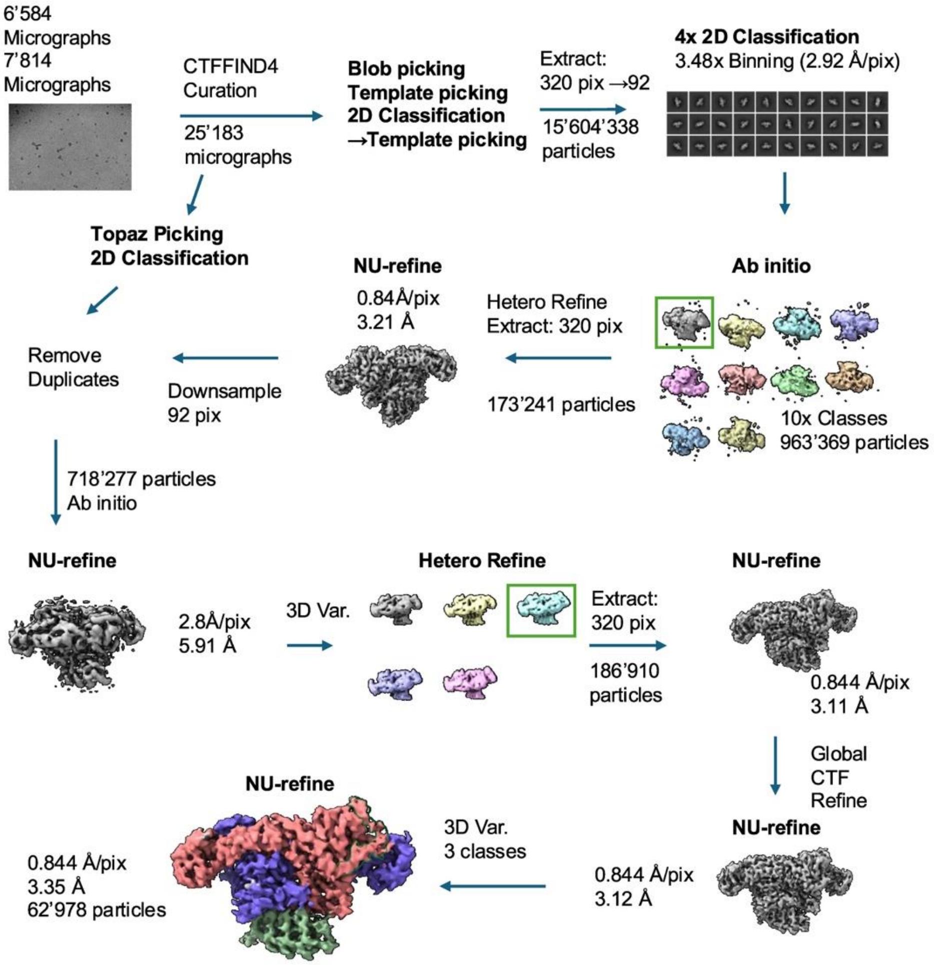

**Fig. S6: CryoEM data processing workflow for GephFL dimer of dimers.**

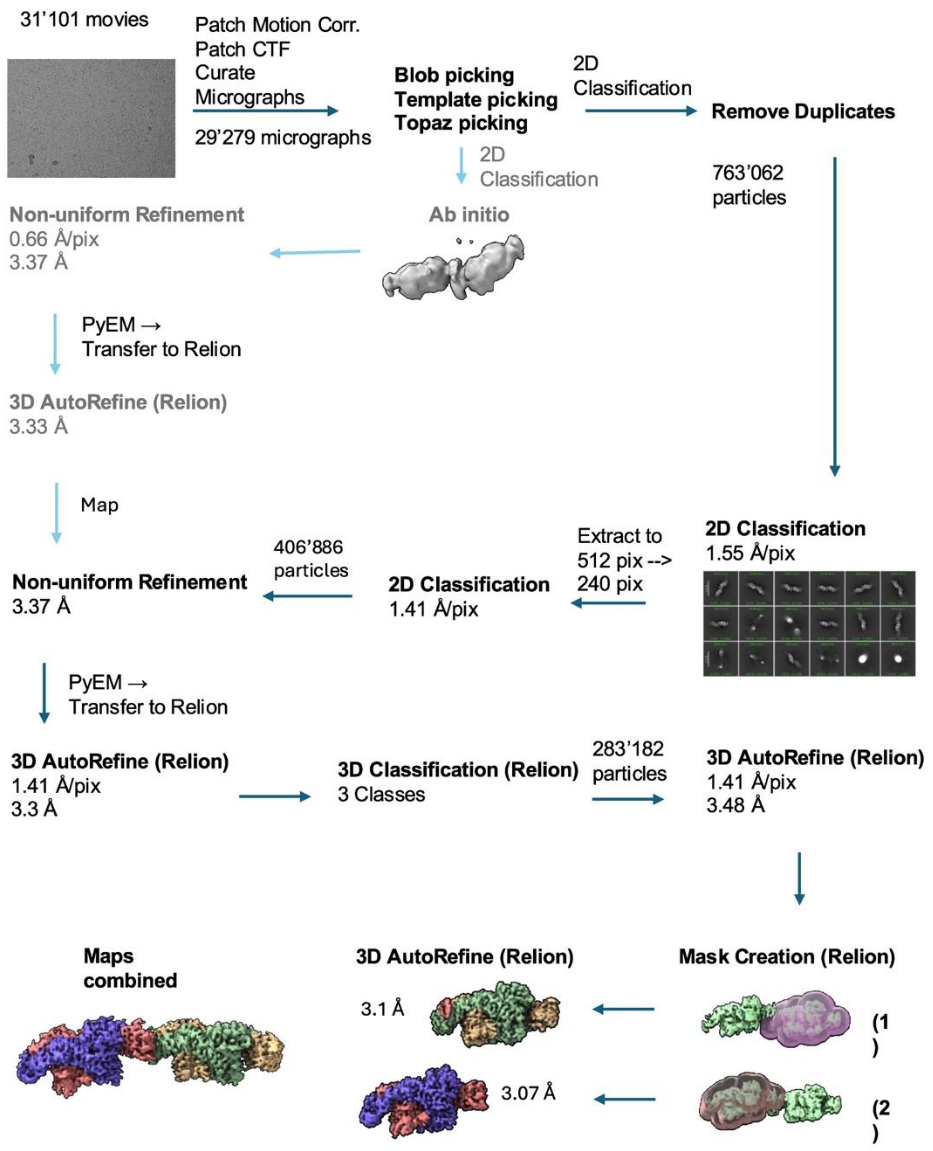

#### Fig. S7: Condensate formation of gephyrin wild-type and variants in HEK293T cells.

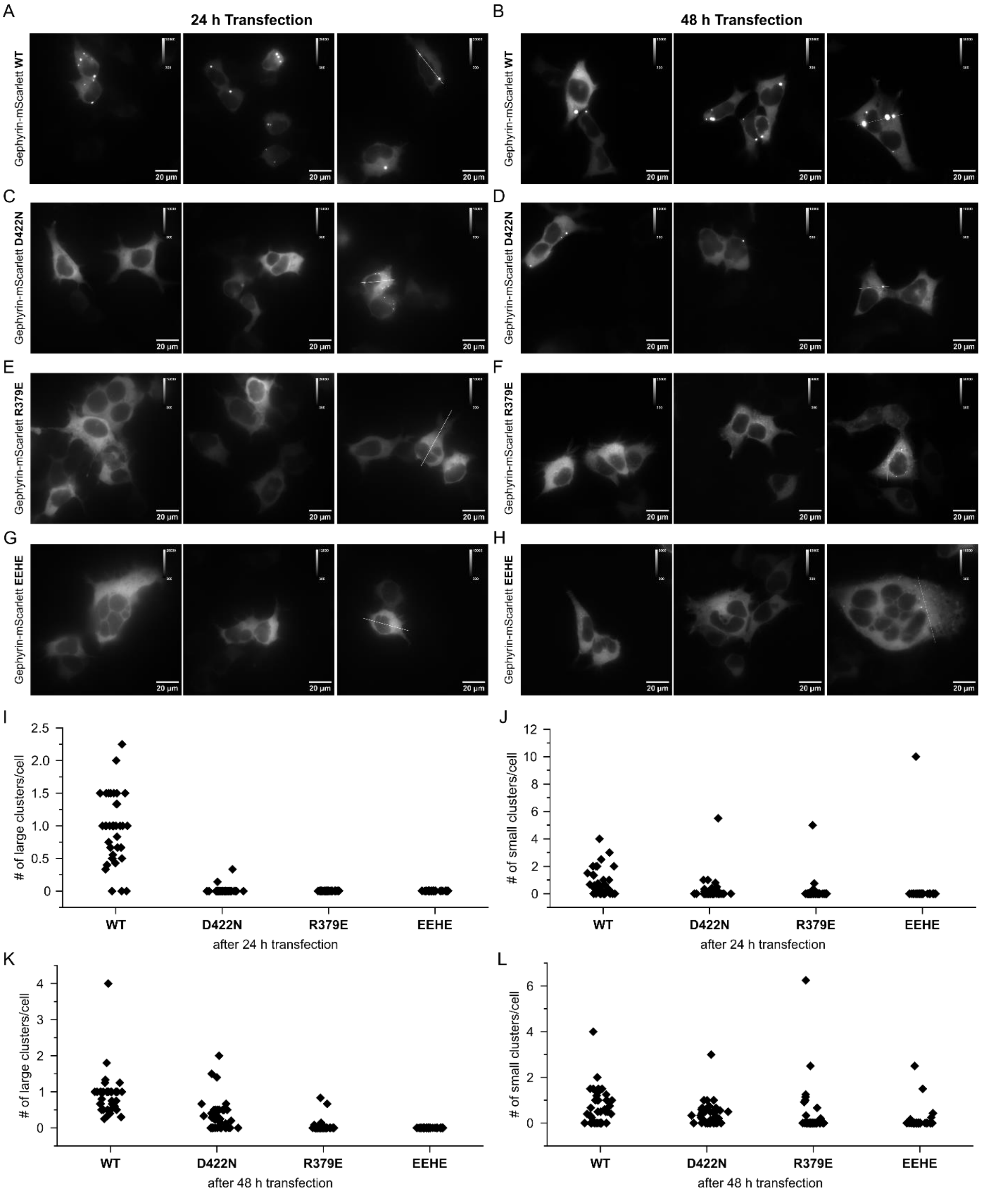


**a-h** Representative wide-field fluorescence microscopy images of fixed HEK293 cells overexpressing mScarlett-labeled gephyrin WT (**a**, **b**), D422N (**c**, **d**), R379E (**e**, **f**) and $^{314}$EEHE$^{317}$ (**g**, **h**) variants. Cells were either transfected for 24 h (**a**, **c**, **e**, **g)** or 48 h (**b**, **d**, **f**, **h**) prior to fixation. **i-j** Cluster counts per cell for the individual mutants. Cells were transfected for 24 h prior to fixation.

Clusters with intensity values 5000-25000 were identified as small clusters (left), ≥25000 as large clusters (right), n≥40. **k-l** Cluster counts per cell for the individual mutants. Cells were transfected for 48 h prior to fixation. Clusters with intensity values 20000-50000 were identified as small clusters (left), ≥50000 as large clusters (right), n≥40.

**Fig. S8. Chromatography of the GephFL P2 variant.**

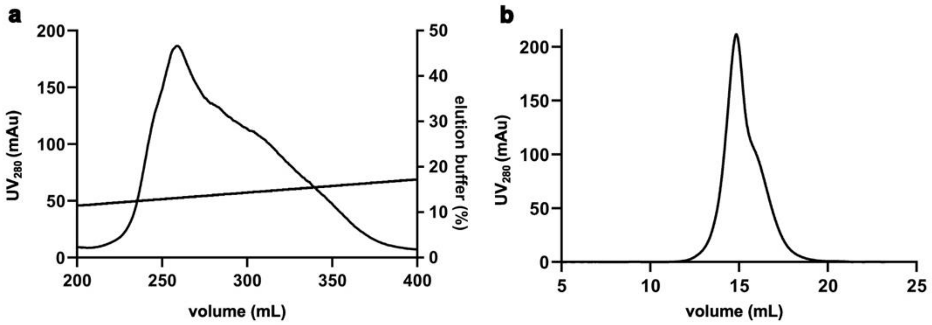


**a** Anion exchange chromatogram of the P2 variant. In contrast, to the chromatogram of the P1 variant with two peaks and a trailing shoulder (Fig. 1b) a main peak with a broad shoulder (representing degradation products) is observed. **b** Size-exclusion chromatogram of the P2 variant using the main peak of (**a**). A main peak with a trailing shoulder is observed instead of the multiple oligomeric species obtained with the P1 variant (Fig. 1c).
